# Study of the hepatotoxicity of trans-coric acid isolated from *Scutellaria baicalensis* in *in vivo* experiments

**DOI:** 10.1101/2024.04.07.588492

**Authors:** Fedorova Anastasia Mikhailovna, Vesnina Anna Dmitrievna, Milentyeva Irina Sergeevna, Prosekov Alexander Yuryevich

## Abstract

Biologically active supplements (BAS) based on plant material, possessing a wide range of beneficial properties (antioxidant, anti-inflammatory, antimicrobial, etc.), are becoming increasingly popular worldwide. Such supplements are readily available and can be used without consulting a healthcare professional. Consequently, the extent of their use is underestimated by doctors. Plant hepatotoxicity is not uncommon, but the frequency and manifestation are not sufficiently characterized. This study is dedicated to investigating the hepatotoxicity of trans-coric acid isolated from the root culture of *Scutellaria baicalensis* in *in vivo* experiments. This work aims to evaluate the hepatotoxicity of trans-coric acid on male rats at doses of 50.0 and 100.0 mg/kg. During the experiment, it was found that trans-coric acid, isolated from the root culture extract of *Scutellaria baicalensis*, does not induce hepatotoxicity in the presented doses in model animals with 14-day intragastric administration. Thus, this study demonstrates that trans-coric acid can be safely used as a component of BAS for the prevention of chronic diseases, thereby promoting healthy aging.

## Introduction

Adhering to a proper diet (consumption of fruits, vegetables, and various antioxidant-rich, anti-inflammatory, and other bioactive substances (BAS)) contributes to improving human health and healthy aging. However, due to economic, climatic, geographical, and other limitations, the systematic consumption of certain products is difficult [1, 2]. Therefore, the development of biologically active supplements (BAS) is relevant.

According to research data, approximately 50% of the adult population in developed countries take BAS containing plant-derived bioactive substances [3]. However, many people are insufficiently informed about the active components of dietary supplements, their side effects, and interactions with medications. There are numerous health risks associated with uncontrolled intake of dietary supplements: heavy metal content, bacterial and fungal infections, risks leading to the development of oncological or chronic diseases [4]. Currently, one-fifth of cases of chemical liver damage are caused by uncontrolled intake of preparations and dietary supplements based on plant materials by specialists, thus increasing the relevance of this problem. Liver damage associated with the consumption of dietary supplements accounts for 2–16% of all identified cases of hepatotoxicity [5]. Toxic liver injury (hepatotoxicity) is a particular type of inflammation and/or structural changes in the organ, often leading to the death of liver cells, caused by medicinal drugs, herbs, and dietary supplements based on them [6]. Such liver damage is often recognized late, as their negative effects are considered minimal, and specific blood tests to exclude causes are currently unavailable.

In Asian countries, where the consumption of dietary supplements is quite popular, the highest prevalence of liver injuries associated with the consumption of dietary supplements is considered: 72% in Korea, 70% in Singapore, and 43% in China [7, 8, 9].

The frequency of occurrence of plant hepatotoxicity is largely unknown. The clinical presentation and severity can be highly variable, ranging from mild hepatitis to acute liver failure requiring transplantation. Scoring systems for assessing the causality of drug-induced liver injury may be useful, but they have not been validated for herb-induced hepatotoxicity. The hepatotoxicity features of widely used herbal products, such as Ayurvedic (*Psoralea corylifolia L*., *Centella asiatica, Atractylis gummifera, Callilepsis laureola* (Impila), *Larrea tridentate*) and Chinese (*Lypocodium serratum; Ephedra sinica; Shosaikoto, Dachaihutang, Xiaochaihutang; Gardenia jasminoides; Paeonia spp*., *Polygonum multiflorum*) herbs, germander flowers (*Teucrium chamaedrys*), greater celandine (*Chelidonium majus*), green tea (*Camellia sinensis*), Herbalife products (Los Angeles, California, USA), Hydroxycut, kava (*Piper methysticum*), peppermint oil, pyrrolizidine alkaloids (*Senecio, Heliotropium, Crotalaria, and Symphytium* (comfrey)), have been individually reviewed by scientists K. Bunchorntavakul and K. R. Reddy [10]. In addition, numerous other plant products, including camphor oil (*Cinnamomum camphora, Vicks VapoRub*), kavakava, saw palmetto, noni juice (*Morinda citrifolia*), cascara (*Cascara sagrada*), mistletoe (*Viscus album*), skullcap (*Scutellaria*), valerian (*Valeriana officinalis*), senna (*Cassia angustifolia* and *C. acutifolia*), usnic acid, neem oil (*Azadirachta indica*), as well as dietary supplements including vitamin A and linoleic acid, can cause hepatotoxicity [Hepatotoxicity associated with herbal and botanical dietary supplements [11, 12].

The main issue is that BAS are not subjected to the same marketing safety or efficacy requirements as pharmaceutical drugs [13]. Since there is no diagnostic biomarker, there are limitations in assessing the causality of hepatotoxicity. Diagnosis of hepatotoxicity is hindered by the fact that patients usually do not perceive BAS as dangerous and therefore do not always inform their doctor about BAS intake. It should be noted that BAS intake may not always be regular. Additionally, the ingredients in BAS can vary significantly and may not always be adequately reflected on the product label. Currently, many literature data point to an important problem associated with BAS consumption [14, 15, 16]. Joint efforts of scientists, physicians, and health authorities are necessary to identify new cases of hepatotoxicity and to educate the public about the risks of BAS consumption.

In this regard, the goal is to study the hepatotoxicity of trans-coric acid, actively included in BAS, isolated from the root culture of *Scutellaria baicalensis* in *in vivo* experiments

In previous studies by the authors, it was established that metabolites isolated from the root culture of *Scutellaria baicalensis* did not cause death or toxic effects on *Caenorhabditis elegans* [17]. In this work, rodents are chosen as model organisms, which are more complex and similar to humans [18, 19].

### Objects and methods of research

The object of the research is trans-coric acid obtained from the water-alcohol extract of *in vitro* root culture of *Scutellaria baicalensis*.

Trans-coric acid extracted from the root culture extract of *Scutellaria baicalensis* was obtained at earlier stages. The methods of cultivating the root culture of *Scutellaria baicalensis*, the process of extraction, extraction, and purification of trans-coric acid are described in the work of A.M. Fedorova and her colleagues [19]. For the transformation of seedlings grown for 14–28 days on a nutrient medium with the following composition: macro salts B5 – 50.00 mg, micro salts B5 – 10.00 mg, Fe-EDTA – 5.00 ml; thiamine – 10.00 mg; pyridoxine – 1.00 mg; nicotinic acid – 1.00 mg; sucrose – 30.00 g; inositol – 100.00 mg; 6-benzylaminopurine – 0.05 mg; indoleacetic acid – 1.00 mg; agar – 20.00 g, strains of soil agrobacteria *Agrobacterium rhizogenes* 15834 Swiss (Moscow, Russia) were used. For in vitro root cultures of *Scutellaria baicalensis*, the cultivation cycle was 5 weeks. The parameters for obtaining the water-alcohol extract of the root culture of *Scutellaria baicalensis* were 30±0.2% ethanol in a ratio of raw material/extractant 1:86 at a temperature of 70±0.1 °C for 6±0.1 hours. The scheme of extraction and purification of trans-coric acid obtained from the root culture extract of *Scutellaria baicalensis* is presented in Figure 1. The degree of purification of trans-coric acid was not less than 95%.

**Figure 1.**
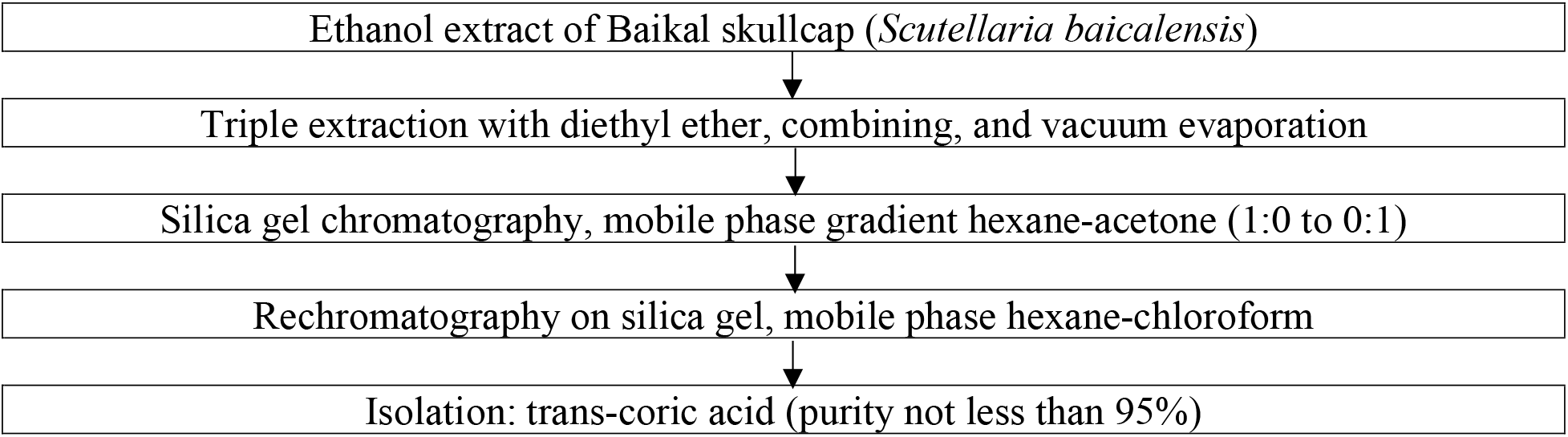
Scheme of purification of trans-coric acid obtained from the root culture extract of *Scutellaria baicalensis*

The *in vivo* studies were conducted at LLC “Iphar”. The use of animals in this study and the conditions of their maintenance during the research were reviewed by the organization’s bioethical committee to ensure compliance with the Policy for Laboratory Animal Care of LLC “Iphar” and regulatory documents governing work with laboratory animals [20–25].

To evaluate hepatotoxicity, 15 male rats of *Rattus sp*. were used, with a health status: free from pathogenic microflora, age at the beginning of the study – 28 weeks, body mass range of animals at the beginning of the study – 623–755 g.

Male animals were chosen because males do not have an estrous cycle, upon which the susceptibility to etiological factors in females depends. Conducting the study on males is therefore preferable [20].

To study the hepatotoxic properties, blood was collected from the jugular vein of the animals, from which serum was obtained and the following indicators were determined: alanine aminotransferase (ALT) activity, aspartate aminotransferase (AST), gamma-glutamyltransferase (GGT), alkaline phosphatase (ALP), total protein, albumin, and globulin levels (difference between total protein and albumin concentration). Measurements were performed on a biochemical analyzer (Minitecno LIND 126 I.S.E. S.r.l, Italy) using commercial kits (Vector-Best Company, Novosibirsk).

The assessment of hepatotoxicity of trans-coric acid was conducted on male rats at doses of 50.0 mg/kg and 100.0 mg/kg administered intragastrically daily for 14.0 days. Purified freshly prepared water was used as the negative control substance and carrier for test substances. The study design is presented in Table 1.

**Table 1.** Design of the hepatotoxicity study

| Group № | Number of animals in the group | Substance given, dose |
| --- | --- | --- |
| 1 | 5 | Purified water |
| 2 | 5 | 50,0 mg/kg |
| 3 | 5 | 100,0 mg/kg |

Daily observations were made of the general condition of the animals, and once a week their body weight and fecal consistency were evaluated. Animals deprived of food the day before were euthanized 24 hours after the last administration. To assess potential liver toxic damage, histological analysis of the liver was performed, the liver weight coefficient (percentage ratio of organ weight to body weight) was determined, and markers of hepatocyte and bile duct damage were determined in serum: ALT, AST, GGT, ALP, total protein, and albumin on a semi-automatic biochemical analyzer. The concentration of globulins was determined as the difference between the concentrations of total protein and albumin.

Since there were no more than 5 animals in each group, the Mann-Whitney test, a non-parametric statistical method, was used to compare the indicators between different groups as it is most suitable for such small samples. Outliers were assessed using the Grubbs test. Differences were considered statistically significant at p < 0.05. The results in the final report are presented using the mean value of the feature X and the standard error of the mean SE.

## Results and Discussion

The body weight data of male rats during the period of intragastric injection of the test substance are presented in Table 2.

**Table 2.**
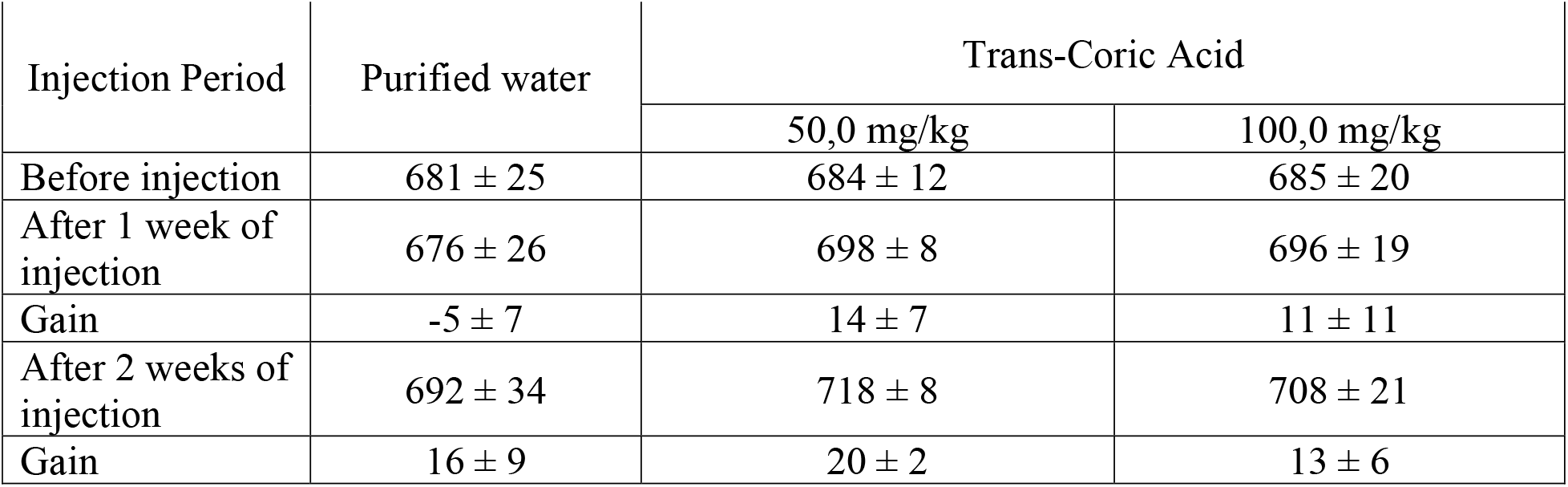
Average body weight data of animals during the period of intragastric injection of trans-coric acid

During the administration period, the animals receiving trans-coric acid did not show any difference in body weight gain dynamics compared to the control animals (p > 0.05).

It was found that trans-coric acid, when injected intragastrically for 14 days at doses of 50.0–100.0 mg/kg, did not affect the body weight of rats, indicating no impact on their overall condition.

Individual blood biochemistry parameters of rats at the end of the intragastric injection period of the test substances are presented in Figure 1.

**Figure 1.**
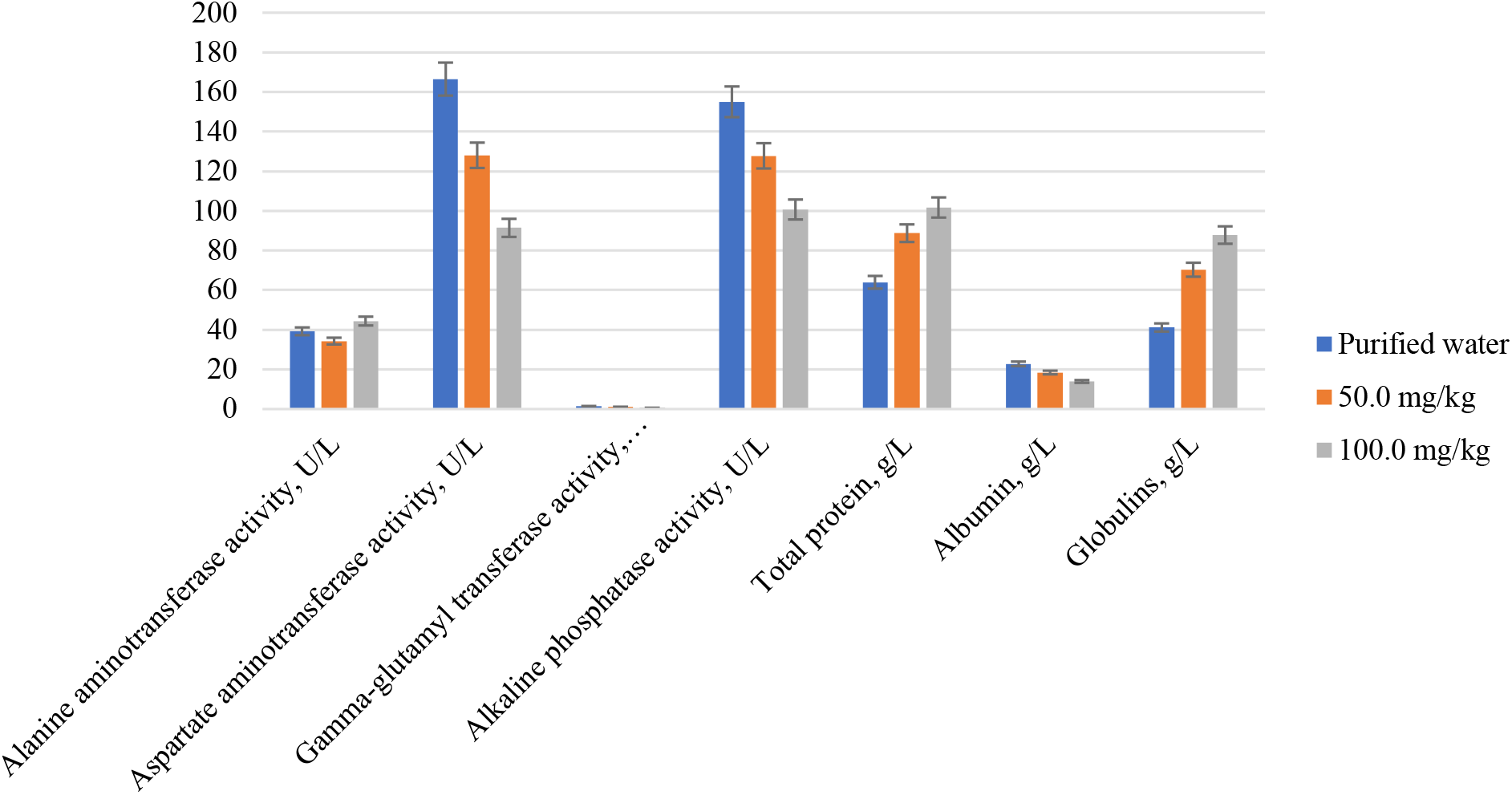
Average blood biochemistry parameters of animals at the end of the period of intragastric administration of trans-cinnamic acid.

The results of blood biochemistry analysis in animals showed changes in the quantitative composition of total protein in serum, as well as a decrease in albumin concentration and a twofold increase in globulins (statistically significant in animals receiving trans-cinnamic acid at a dose of 100.0 mg/kg, and not statistically significant at a dose of 50.0 mg/kg). Administration of trans-cinnamic acid at a dose of 100.0 mg/kg also led to a statistically significant decrease in AST, GGT, and ALP activity. A characteristic biochemical sign of hepatotoxicity is a significant increase in the activity of liver enzymes (primarily ALT and AST). Therefore, the observed decrease in liver enzyme activity in the experiment cannot be interpreted as a result of the hepatotoxicity of the test substances, except in cases of critical liver damage (which was not observed in this experiment). Thus, the decrease in GGT, AST, and ALP activity does not indicate the hepatotoxicity of trans-cinnamic acid.

The change in the composition of total serum protein in animals under the influence of trans-cinnamic acid is a manifestation of the specific pharmacological activity of this substance, rather than a consequence of chronic inflammatory processes [23].

Therefore, based on the obtained data, trans-cinnamic acid, when administered intragastrically for 14 days at doses of 50.0–100.0 mg/kg, does not cause changes in blood biochemistry indicative of hepatotoxicity of these test substances.

No pathological changes were detected during the planned necropsy. Thus, based on the obtained data, trans-cinnamic acid, when administered intragastrically for 14 days at doses of 50.0–100.0 mg/kg, does not cause pathological changes in the liver.

The indicators of absolute and relative liver mass of animals at the end of the period of intragastric administration of trans-cinnamic acid are presented in Table 3.

**Table 3.**
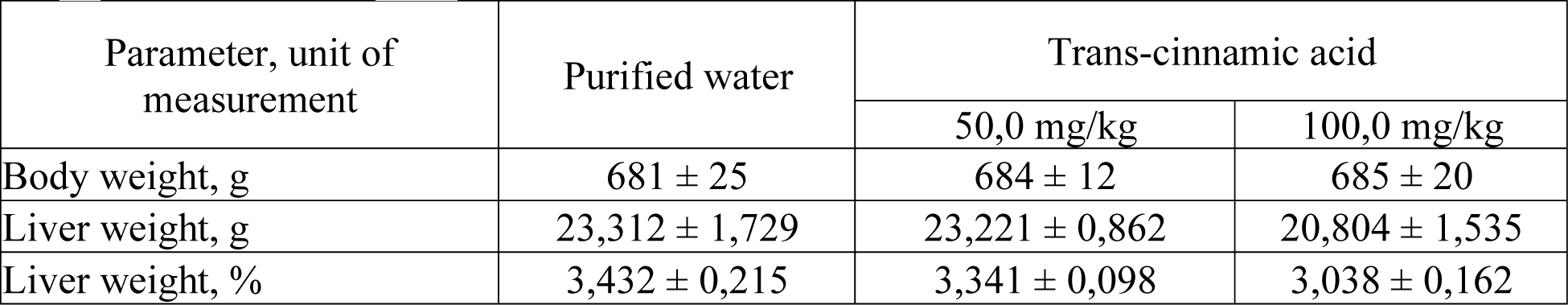
Average values of absolute and relative liver mass of animals at the end of the intragastric administration period of trans-cinnamic acid

Statistically significant differences in the body weight and liver mass between intact animals and animals receiving the test substances were not detected.

Thus, according to the obtained data, trans-cinnamic acid, when administered intragastrically for 14 days at doses of 50.0–100.0 mg/kg, does not induce pathological changes in the liver.

Microscopic analysis of liver structures in all experimental animals revealed congestion with erythrocyte stasis, stromal edema, and focal granular or hydropic hepatocellular degeneration. These phenomena were observed in all groups, including the control group, and are attributed to acute circulatory disturbance at euthanasia.

Therefore, based on the obtained data, trans-cinnamic acid, when administered intragastrically for 14 days at doses of 50.0–100.0 mg/kg, does not induce any morphological changes in liver tissues.

It was found that the observed decrease in the activity of gamma-glutamyl transferase, aspartate aminotransferase, and alkaline phosphatase does not indicate hepatotoxic activity of trans-cinnamic acid, as a significant increase in the activity of liver enzymes (especially alanine aminotransferase and aspartate aminotransferase) is a characteristic biochemical sign of hepatotoxicity, which was not observed. Therefore, trans-cinnamic acid, when administered intragastrically for 14 days at doses of 50.0–100.0 mg/kg, does not induce changes in blood biochemistry indicative of hepatotoxicity of these test substances.

## Conclusion

This study aimed to investigate the hepatotoxic properties of trans-coric acid extracted from the root culture of *Scutellaria baicalensis* in *in vivo* experiments. During the course of the study, it was found that the administration of trans-coric acid did not cause severe health disorders or mortality in the model animals - *Rattus sp*. Throughout the entire administration period, there were no signs of health impairment in any of the animals related to the toxic effects of trans-coric acid, including changes in fecal consistency. Thus, based on the obtained data, trans-coric acid at doses of 50.0–100.0 mg/kg injected intragastrically for 14 days did not cause significant signs of health impairment. Therefore, supplements containing these doses of trans-coric acid are relatively safe and advisable for consumption.

## Funding

*The work was carried out as part of the state assignment on the topic* “*Development of biologically active additives consisting of metabolites of plant objects in vitro, for the protection of the population from premature aging” (project FZSR-2024-0008)*.

*The work was conducted using the equipment of the Centre for Collective Use* “*Instrumental methods of analysis in the field of applied biotechnology” at KemSU*.

